# An Easy-to-Fabricate Cell Stretcher Reveals Density-Dependent Mechanical Regulation of Collective Cell Movements in Epithelia

**DOI:** 10.1101/2020.08.24.265629

**Authors:** Kevin C. Hart, Joo Yong Sim, Matthew A. Hopcroft, Daniel J. Cohen, Jiongyi Tan, W. James Nelson, Beth L. Pruitt

## Abstract

**Introduction:** Mechanical forces regulate many facets of cell and tissue biology. Studying the effects of forces on cells requires real-time observations of single- and multi-cell dynamics in tissue models during controlled external mechanical input. Many of the existing devices used to conduct these studies are costly and complicated to fabricate, which reduces the availability of these devices to many laboratories.

**Methods:** We show how to fabricate a simple, low-cost, uniaxial stretching device, with readily available materials and instruments that is compatible with high-resolution time-lapse microscopy of adherent cell monolayers. In addition, we show how to construct a pressure controller that induces a repeatable degree of stretch in monolayers, as well as a custom MATLAB code to quantify individual cell strains.

**Results:** As an application note using this device, we show that uniaxial stretch slows down cellular movements in a mammalian epithelial monolayer in a cell density-dependent manner. We demonstrate that the effect on cell movement involves the relocalization of myosin downstream of Rho-associated protein kinase (ROCK).

**Conclusions:** This mechanical device provides a platform for broader involvement of engineers and biologists in this important area of cell and tissue biology. We used this device to demonstrate the mechanical regulation of collective cell movements in epithelia.

## Introduction

Mechanical force regulates many cellular functions underlying tissue morphogenesis, including differentiation, proliferation, and collective cell migration.^6,7,8,12,13,25,26,28,34,38^ To study effects of mechanical forces on cells and tissues, a number of devices have been developed that apply either tensile or compressive strains to cells.^10,11,14,15,16,18,22,29,36,37,41,^ However, the fabrication of these devices often requires custom engineering design and access to sophisticated facilities such as a clean room or machine shop, which are not readily available to most laboratories. Moreover, many of these devices are incompatible with high-resolution imaging modalities that are indispensable to explore how cells transduce mechanical inputs into downstream signaling outputs. Our goal, therefore, was to build an easy-to-fabricate, low-cost, uniaxial mechanical stretching system compatible with high-resolution, fluorescence time-lapse microscopy. We achieved our goal by using low-cost 3D-printed molds, optically clear polydimethylsiloxane (PDMS) components, and a vacuum-driven stretching design.

Mechanical devices require not only electronic control of the amount of force and the timing of force application, but also quantification of resulting cell strain. Thus, we also constructed a pressure controller that induces a repeatable degree of stretch to monolayers, and developed a custom MATLAB code, called CSI (<u>C</u>ell <u>S</u>train from <u>I</u>mages), to estimate individual cell strains. Typically, whole-field strain levels are calibrated using fiducial objects, e.g., microspheres and patterned fluorescent proteins placed on the stretching substrates, or following distinguishable membrane and cell defects, ^16,23,29^ and then assuming a uniform strain across the attached cells during an experiment. However, such approaches do not directly measure and map individual cell strains. Our quantification method allows a direct measurement of cell strain mapped onto an image of individual cells in the experiment.

To showcase the features of our device, we used transmitted-light time-lapse microscopy to follow the movement of cells within an epithelial monolayer under mechanical stretch. We show that cellular movement initially slowed down, but the average speed eventually returned to levels observed before the application of stretch. Finally, we used epifluorescence microscopy to show that the change in cell movement speed depends on density-dependent myosin activation downstream of Rho-associated protein kinase (ROCK).

## Materials and Methods

### Uniaxial stretch device design

We designed an easy-to-fabricate, pressure-regulated device compatible with live-cell imaging. Our uniaxial cell stretching device consists of two layers of flexible PDMS (Figure 1A).^35^ The bottom layer comprises a 125 μm thick pre-fabricated silicone membrane (SMI Manufacturing) that serves as the cell-culture substrate. The top layer features a cell-culture chamber with thin, deformable walls that separate it from the surrounding vacuum chamber compartment. Vacuum is applied to the pneumatic chamber, which deflects the sidewalls outward from the cell-culture chamber and applies mechanical strain across the short axis of the cell-culture membrane (Figure 1A). The stretch is uniaxial due to the high aspect ratio of the cell-culture chamber. The overall device is the same size as a standard microscope slide (75 mm × 25 mm), and fits on a standard microscope stage for easy use with most commercial imaging systems (Figure S1).

**Figure 1:**
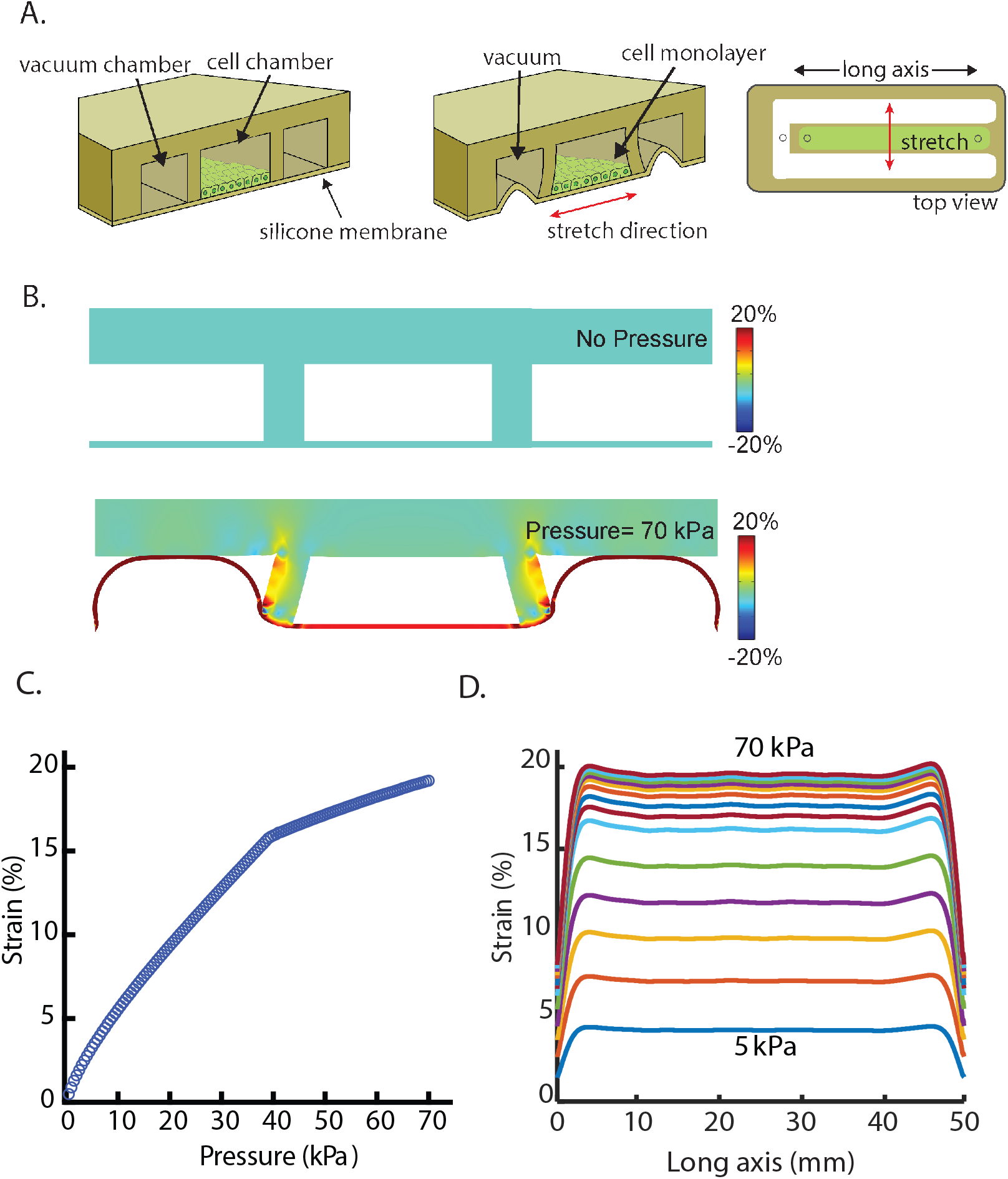
Design and computational analysis of a low-cost, easy-to-fabricate, pneumatically controlled uniaxial cell stretching device. A. Schematic illustrations of the uniaxial stretch device in the cut-away side view before stretching (left) and after stretching (center) and the top view of the device (right). When vacuum pressure is applied to the two side vacuum chambers, the side chamber walls are deflected outward from the cell-culture chamber, resulting in the suspended silicone membrane being stretched. The stretching direction is perpendicular to the long axis of the cell-culture chamber. B. Finite element analysis (FEA) example of the uniaxial stretch device before and after application of vacuum pressure to the side chambers. The color intensity indicates nodal strain calculated in the lateral stretch direction. Without applying vacuum pressure, no strain is applied to the membrane in the cell-culture chamber (top). Upon applying a vacuum pressure of 70 kPa, the cell-culture membrane is predicted to undergo 19% strain. (bottom). C. FEA prediction of the strain profile of the cell-culture membrane corresponding to the pressure applied to the vacuum chamber. At 37 kPa, the membrane of the vacuum chamber makes contact with the top of the vacuum chamber, modeled as a contact event in the FEA model. D. FEA prediction of the strain profile of the cell-culture membrane along the long axis of the device with applications of vacuum pressures from 5 kPa to 70 kPa, demonstrating the homogeneity of strain. Every 5 kPa is depicted with a line that follows the strain (%) of the device along the long axis of the device.

### Fabrication of the stretch device

Figure 2 outlines the steps involved in the fabrication of the device. A mold for the PDMS top layer is cast by 3D-printing using a commercial 3D printer (Dimension 1200es BTS, Stratasys) and the thermoplastic polymer, acrylonitrile butadiene styrene. A Solidworks (Dassault Systems) design file for the mold is available for download from the Open Science Framework archival Project Repository (https://osf.io/gtrju/). Sylgard 184 PDMS (Dow Corning, Inc.) at a ratio of 10:1 (elastomer: curing agent) is poured onto the mold, degassed for 1 hr in a vacuum chamber to minimize trapped air bubbles and improve device clarity, and then cured in a 65°C oven for 4 hr on a level shelf, or on a leveled hotplate at 65°C. The PDMS is peeled off the mold, and inlet and outlet holes for cell loading and vacuum are punched using a 1 mm diameter biopsy punch (Acuderm Acu-Punch, Fisher Scientific). To remove roughness on edges of the PDMS from the 3D printed mold, the uncured PDMS is placed on a thin sheet of uncured PDMS, and then moved onto a clean surface and baked in a 65°C oven for 4 hr on a level shelf. After cleaning the PDMS surface with adhesive tape (Scotch tape, 3M), the device and the prefabricated silicone membrane are plasma-treated for 45 sec at 0.1 Torr and 18 W (PDC-32G, Harrick Plasma). Alternatively, a handheld plasma wand can be used for this step. Finally, the device is bonded to the membrane using light hand pressure for 10 seconds, and excess material trimmed to the edge of the slide using a single-edged razor blade.

**Figure 2:**
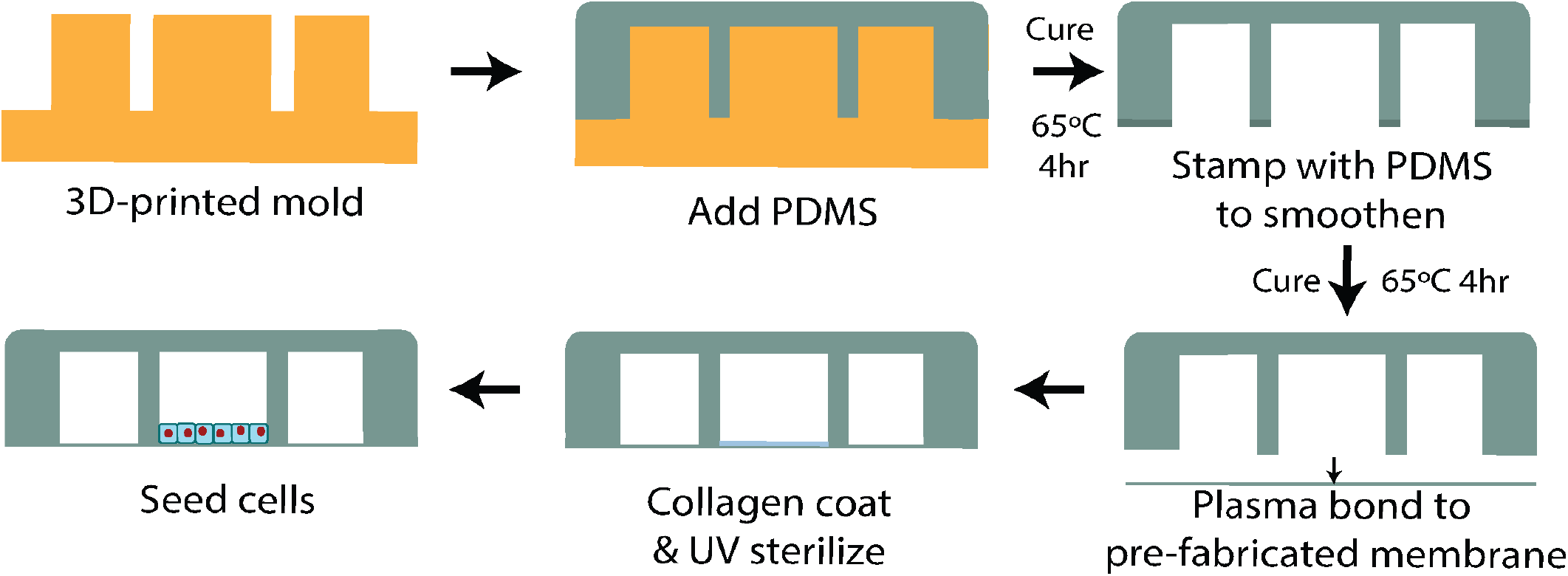
Representative side views of the step-by-step fabrication of the uniaxial stretch device and preparation for cell culture. A PDMS layer with a cell-culture chamber and two vacuum chambers is created by using a 3D-printed mold. After removing the roughness on edges of the PDMS from the 3D-printed mold, the PDMS layer is plasma bonded with the prefabricated silicone membrane. After the cell-culture chamber is coated with extracellular matrix and UV-sterilized, cells are seeded and grown as a confluent monolayer.

### Finite Element Analysis of Cell Stretching Device

A 2D finite element analysis (FEA) model was made using COMSOL Multiphysics software (version 4.4, COMSOL Inc.) that included the suspended membrane, a sidewall, and a top layer with a half-symmetric boundary condition for the cross-sectional area of the cell loading chamber. The FEA model is available for download from the Open Science Framework archival Project Repository (https://osf.io/gtrju/). To predict the change in strain when the bottom membrane contacts the top layer as the vacuum increases, we defined a contact pair between the top layer (destination boundary) and the bottom membrane (source boundary). Assuming that gravity has negligible effects, the model was simplified by applying pressure on the area around the vacuum chamber at increments of 1 kPa, from 0 kPa to 70 kPa. The PDMS material of the device was assumed hyperelastic (Mooney–Rivlin model) and isotropic, and the silicone membrane was assumed to have the same mechanical properties as the PDMS. The mechanical properties of PDMS were assumed to be as follows: a modulus of elasticity of 1 MPa, a Poisson’s ratio of 0.49, and a density of 1000 kg/m^3^.^20^ To predict the strain distribution of the cell loading chamber along the long axis, an additional 3D FEA model was implemented with a half-symmetric boundary condition at the cross-sectional area in the half of short axis (i.e., x axis) of the cell loading chamber and 46,458 nodes of tetrahedral meshes for the bottom membrane and the sidewalls. Parametric analysis was conducted to predict strain increases with applied pressure until there was contact between the two layers between 0 kPa and 35 kPa at increments of 5 kPa. The model was simplified without contact mechanics between the bottom membrane and the top layer. The strain levels results from the 3D FEA at 40–70 kPa were assumed linearly proportional to the strain level predicted by the 2D FEA to compensate for the effect of contact between the bottom membrane and the top layer.

### Cell Culture

Parental MDCK GII cells, MDCK cells stably expressing E-cadherin-RFP or GFP-myosin IIA, ^21,39^ were used. Cells were cultured at 37°C and 5% CO_2_ in low-glucose DMEM containing 10% FBS, 1 g/L sodium bicarbonate, and penicillin/streptomycin/kanamycin. The cell-culture chamber was coated with 50 μM type I collagen to facilitate cell attachment and spreading. To coat the cell-culture chamber, collagen I in 0.1% acetic acid (Collagen, rat tail, Type I, Invitrogen) was injected via the inlet of the cell-culture chamber, and the chamber was incubated for 30 min at room temperature (~20°C). Excess collagen solution was flushed out of the chamber with phosphate-buffered saline (PBS pH7.4, Invitrogen), and the device was UV-sterilized for 10 min. A cell suspension (3.5 – 4.0 × 10^5^ cells in a volume of 400 μl) was added to the cell-culture chamber to obtain a low-density, confluent cell monolayer that was ready for imaging 16–20 hr later.

### Cell Imaging with Controlled Stretch

A monolayer of MDCK cells was imaged on the membrane of the device using a customized Zeiss Observer inverted microscope (Intelligent Imaging Innovations, 3I) with 5x or 20x objectives, in a temperature- and CO_2_-controlled incubator. For the quantification of cell strain, cells were imaged at time zero, and re-imaged following increased strain every minute; optical focus was adjusted manually if necessary. The applied strain is controlled by the amount of pressure applied to the device. Typical in-house laboratory vacuum can create a pressure difference of 70 kPa, which creates a maximum strain in the stretcher membrane of ~20%. The pressure difference can be controlled in two ways, depending on the relative availability of: 1) a mechanical pressure regulator (Parker Valve Inc.) that uses a spring balance against the gas flow through the regulator to maintain a constant pressure in the regulated system (Figure S2A); or 2) an electronic pressure controller. Design files and basic code for the controller are available for download from the Open Science Framework archival Project Repository (https://osf.io/gtrju/); assembled controllers with a user interface are available from Red Dog Research (Figure S2B). The electronic pressure controller uses a pressure sensor with 10-bit resolution that operates two valves connected to the vacuum supply line to control the pressure in the cell stretcher. The pressure controller was interfaced with a USB connection and graphical user interface for user input. The controller maintains pressure within 0.25 kPa of the set-point, which is equivalent to 0.1% strain. The controller can execute pre-programmed pressure waveforms, and hence modulate strain in a variety of ways. In this work, a programmed constant pressure was applied to the pneumatic side chambers and used to control the constant strain level of the membrane.

### Cell Strain Analysis and Image Processing

MATLAB (Mathworks, Inc.) code was developed for calculating cellular strains based on the measure of deformation relative to a reference length along the long (axial, ****e_xx_****) and short axes (transverse, ****e_yy_)**** of the cell-culture chamber. The MATLAB code, called CSI, is available for free on Github [https://github.com/MicrosystemsLab/CellStrainImages]. Normal strains ****e_yy_****, ****e_xx_**** and in-plane shear strain ****e_xy_**** are defined in ***x***, ***y*** plane as: ****e_xx_**** = *∂**u_x_**/∂**x**, **e_yy_*** = *∂**u_y_**/∂**y**, **e_xy_*** = (*∂**u_x_**/∂**y** + ∂**u_x_**/∂**x**)/2*, where ****u_x_**** and ****u_y_**** are displacements between particles in the body in ****x, y**** directions. Prior to calculating the strain of each cell, the images were filtered using Gaussian kernel (2 × 2 pixels) and then adjusted for the imbalance of illumination using contrast limited adaptive histogram equalization. To calculate strains, fluorescent cell images were registered before and after applying strain using the ‘imregister’ function of MATLAB with Mattes mutual information similarity metrics. An affine transformation was applied to generate matched images after enlarging, shearing, and translating images in 2-dimensional space. To analyze cell-by-cell strain, cells were segmented using the outlines of plasma membranes delineated by E-cadherin-RFP, and images before and after stretch were compared to calculate the axial, transverse, and shear strains.

### Analysis of cell velocities and myosin localization

Cell monolayers were imaged by phase contrast illumination with a 5x objective every 10 min for 7 hr, and Particle Image Velocimetry (PIV) was used to analyze cell movements.^40^ The PIVlab software in MATLAB was used to analyze a 96 × 48 interrogation window with an overlap of 50%. The mean velocities were averaged from 6 time points over the course of each hour. To calculate normalized speeds, cells were imaged for 1 hr and the mean velocities of cells in the monolayer were calculated, and these velocities were then used to calculate the relative speed for the other time points. To quantify changes in the distribution of myosin-IIA during strain, MDCK cells expressing GFP-myosin IIA were imaged with a 20x objective. ImageJ software was used to quantify the amount of cortical GFP-myosin-IIA in each image as described previously.^13^

### Statistical Analysis

An unpaired Student *t*-test was applied to pairs of treatments for statistical analysis when comparing separate imaging experiments. A paired Student *t*-test was used for statistical analysis for comparing the before and after strain images of the same monolayer of cells.

## Results

### Modeling of predicted membrane strain under stretch

We modeled the mechanical strain of the cell culture membrane of the device under increasing 1 kPa increments of vacuum pressure from 0 kPa to 70 kPa using FEA (Figure 1). The 2D FEA model (COMSOL, Inc.) included contact mechanics to predict and capture the collapse pressure of the membrane. This pressure response captures two regimes: 1) membrane strain during unconstrained stretch; and 2) membrane strain after the bottom membrane in the vacuum chamber contacts the top layer (Figure 1A, B). The model predicted that strain increased linearly with applied pressure up to 37 kPa (Figure 1C, Movie S1), at which point strain increase slowed as the bottom membrane contacted the top of the chamber (Figure 1B). An additional 3D FEA model was created to predict the strain distribution in the cell-culture chamber along the long axis. Figure 1D shows that strain is largely homogeneous along the length of the device at pressures from 5 kPa to 70 kPa due to the high aspect ratio of the cell-loading chamber (50 mm × 4 mm, Figure S1).

### Measuring individual cell strains and shapes under stretch

The strain applied to cells was quantified using CSI, a custom code developed in MATLAB, from real-time images of a monolayer of MDCK cells expressing the cell-cell adhesion protein E-cadherin-RFP. E-cadherin localizes to the plasma membrane and, therefore, outlines the perimeter of each cell. We applied vacuum pressure to the side chambers in increments of 3.4 kPa from 0 kPa to 70 kPa (Movie S2). The CSI code analyzes the acquired images by segmenting and tracking individual cells from the plasma membrane profile of each cell marked by E-cadherin-RFP. We compared each cell before and after stretching to calculate strain in the directions of applied stretch (****e_xx_****), perpendicular to applied stretch (****e_yy_****), and in-plane shear (****e_xy_****). Figure 3A shows cell strains in different directions (at 70 kPa) as color maps within the outline of the cells. Observed strains varied by up to 5%, which we attributed to heterogeneity in the size, shape, and properties of cells due to the low cell density of the cell monolayer. As expected, the strain in the stretching direction (****e_xx_****) increased with applied pressure and exhibited an initial steep increase until 25 kPa, which then plateaued. There were no changes in strain in the direction perpendicular to stretch or in-plane shear (Figure 3A; note that ****e_yy_**** and ****e_xy_**** overlap). Single-cell segmentation allowed us to estimate mechanical stretch-induced changes to cell area, perimeter and eccentricity. Cell area and cell perimeter increased at the same rate as the increase in cell strain (Figure 3B), but there were minimal changes in eccentricity over the range of stretch, indicating that these levels of stretch do not significantly affect cell shape (Figure 3B).

**Figure 3:**
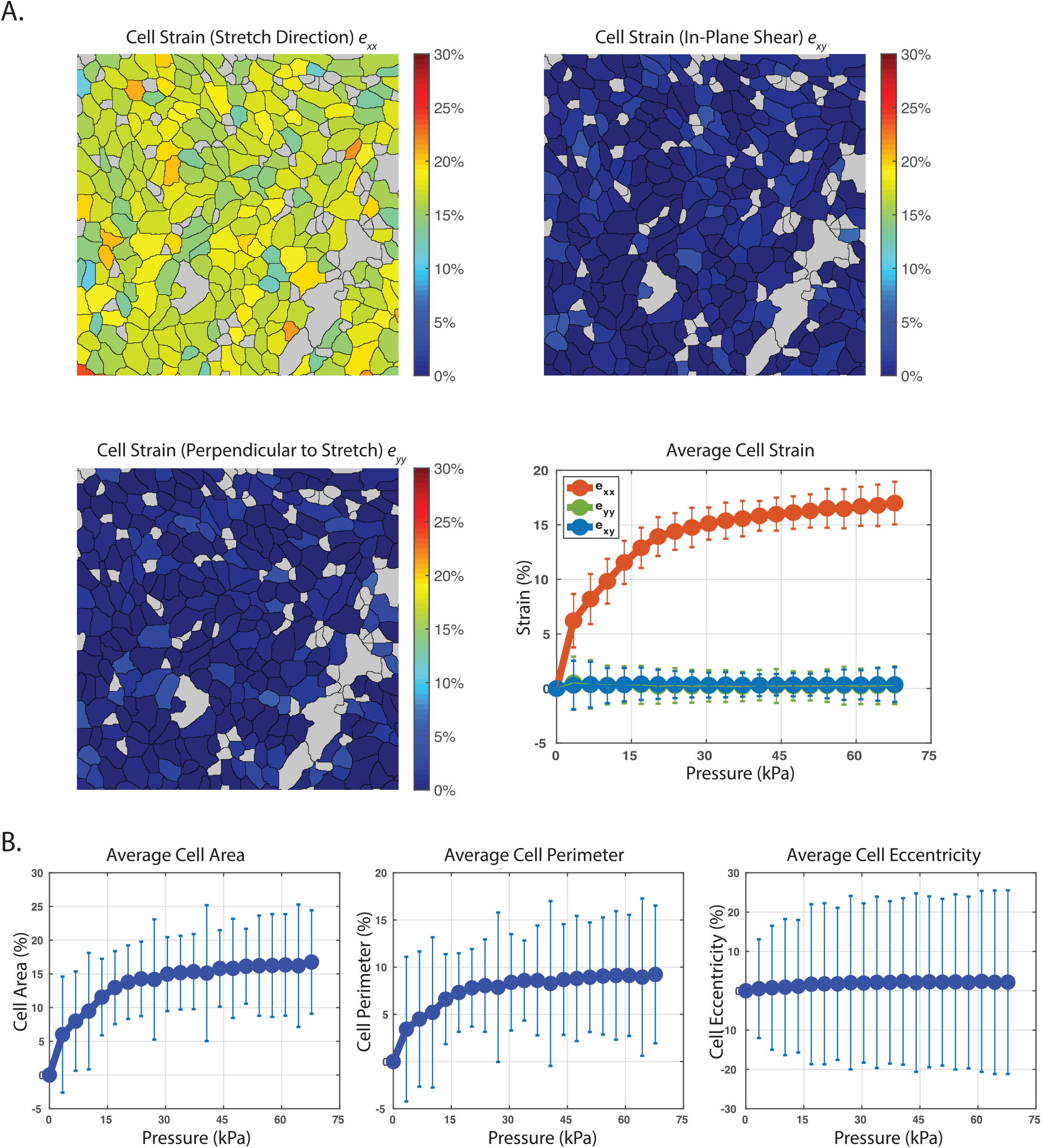
Cell strain analysis and image processing to measure individual cell strain and shape under stretch. A. Cell strain analysis results. First three panels are representative images of cell strain measurements of individual cells in a cell monolayer in the direction of applied stretch (****e_xx_****), perpendicular to applied stretch (****e_yy_****), and in-plane shear (****e_xy_****) at 70 kPa of vacuum pressure. Our custom MATLAB code (CSI) identifies the cell boundaries and measures cell strains at different vacuum pressures. Last panel is the cell strain profile from individual cells at 0–70 kPa in the normal (****e_xx_****), transverse (****e_yy_****), and shear (****e_xy_****) directions from 3 independent experiments. B. Percent changes in average cell area, cell perimeter, and eccentricity (how circular the ellipse is) profile of cells from 3 independent experiments.

### Cell movements slow down in response to stretch

We analyzed the temporal dynamics of collective cell movement within a mechanically stretched MDCK cell monolayer (Figure 4). The cell monolayer was imaged every 10 min for 1 hr with no stretch, and then every 10 min for up to 7 hr during the application of a physiological level of stretch (15%, 40 kPa vacuum).^17^ Using PIV, we generated average velocity maps from these images by averaging the cell speeds for every hour (Figure 4A).^40^ On average, cell movement slowed down by 18.4%, from 11.0 ± 2.9 μm/hr to 9.0 ± 2.2 μm/hr an hour (mean ± SD) during the application of stretch (Figure 4C). In contrast, under no-stretch conditions there was no significant change in average cell velocity during this time (Figure S3). Longer-term tracking of cell movement over 7 hr revealed average cell velocity eventually returned to a level similar to that estimated before the application of stretch (Figure 4D). In contrast, the average velocity of cells not under strain gradually decreased over 7 hr (Figure 4C).

**Figure 4:**
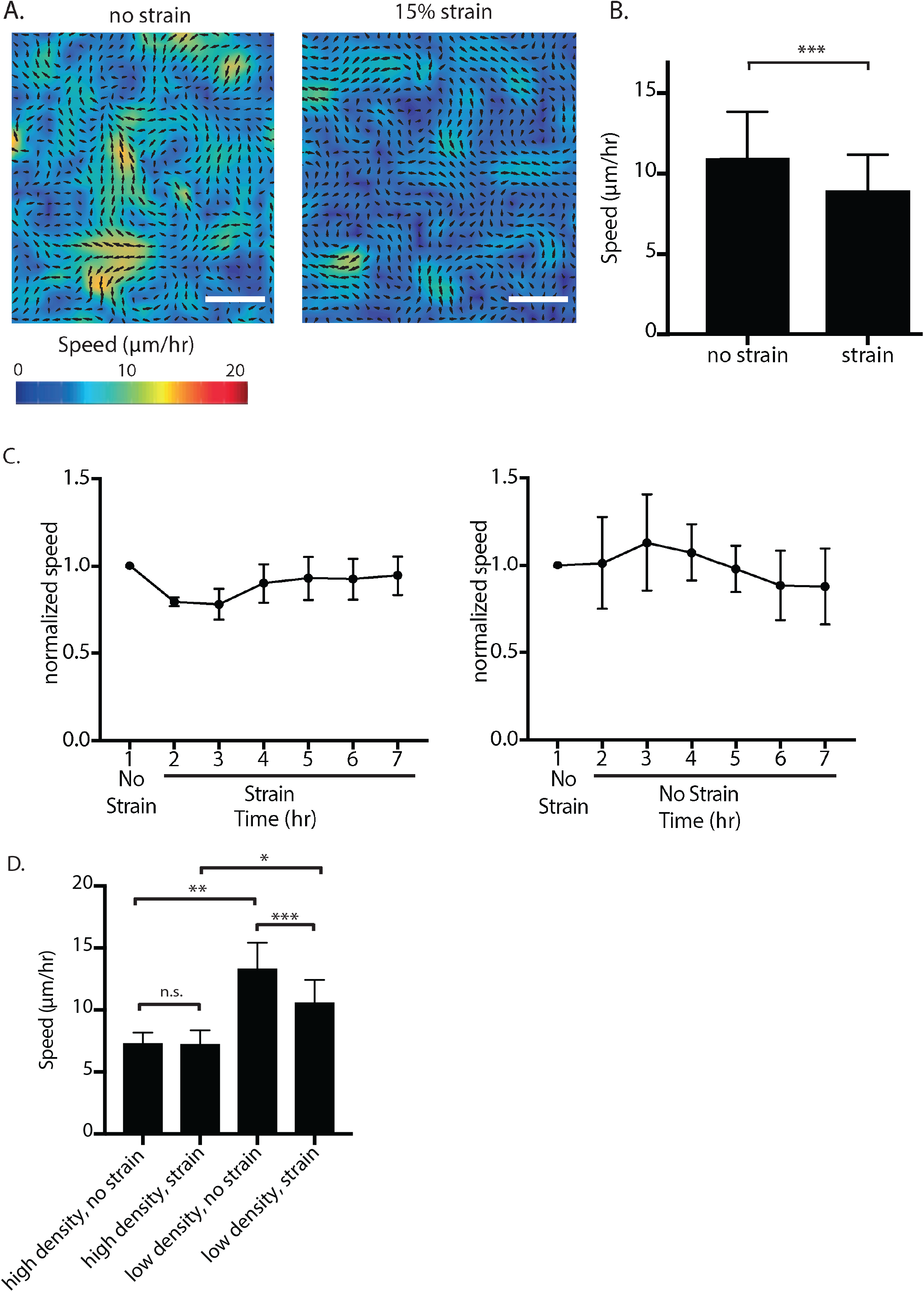
Mechanical strain induces interim slowdown of cell migration in a MDCK cell monolayer. A. Representative mean velocity map from 1 hr of imaging at rest and following 1 hr of imaging with 15% strain. The collective migration velocities were measured with PIV (particle image velocimetry) from phase contrast images taken every 10 min at 5x magnification. Scale bars: 50 μm. B. Mean velocities from 1 hr of no strain followed by 1 hr of 15% strain of the same monolayer. Quantifications were mean +/− SD from 7 independent experiments; paired t-test p values; ***, p<0.001. C. Normalized mean velocities over the course of 7 hr of imaging every 10 minutes with or without strain. Velocities from hours 2-7 were normalized to the first hour of no strain for all experiments. Quantifications were mean +/− SD from 4 independent experiments. D. Mean velocities from 1 hr of no strain followed by 1 hr of 15% strain of the monolayer grown at high density or low density. Quantifications were mean +/− SD from 4 independent experiments; paired t-test p-value for comparison between no strain and strain for the same monolayer (low and high-density); unpaired t-test p-value for comparison between low-density and high-density; ***, p<0.001; **, p<0.01.

In monolayers grown at high density (3.0 ×10^3^/mm^2^ – 4.0 ×10^3^/mm^2^), cells migrated at slower speeds than in lower density (2.0 ×10^3^/mm^2^ – 2.5 ×10^3^/mm^2^) monolayers under no stretch conditions: 7.3 ± 0.9 μm/hr versus 13.3 ± 2.1 μm/hr, respectively (Figure 4D, Figure S4). The average cell velocity in high-density monolayers did not change with the application of mechanical stretch; 7.3 ± 0.9 μm/hr to 7.2 ± 1.2 μm/hr. In contrast, low-density monolayers slowed down within the first hour of applied stretch from 13.3 ± 2.1 μm/hr to 10.5 ± 1.9 μm/hr (Figure 4D, Figure S4).

### Slower cell movements in response to stretch are dependent on cell density and myosin dynamics

Collective cell movements are driven by actomyosin dynamics, and myosin IIA is required for cell migration.^9,30^ We hypothesized that changes in average cell velocity due to mechanical stretch were associated with a change in actomyosin dynamics. MDCK cells expressing GFP-myosin IIA were cultured on the device, and the subcellular localization of GFP-myosin IIA imaged during mechanical stretch (Figure 5A).^21^ Upon application of stretch, the fluorescence intensity of GFP-myosin IIA increased in the cell cortex close to and along the plasma membrane (Figure 5A).^13^ In contrast, the fluorescence intensity of GFP-myosin IIA in cells not under stretch remained diffusely distributed in the cytoplasm, with little accumulation at the cortex or plasma membrane (Figure S5A). Under the same conditions, little or no changes in the distribution of E-cadherin-RFP were detected (Figure S7). The changes in myosin IIA distribution upon stretch was dependent on cell density; accumulation of myosin IIA at the plasma membrane upon stretch was less in high cell density monolayers, than low cell density (Figure 5B, Figure S6).

**Figure 5:**
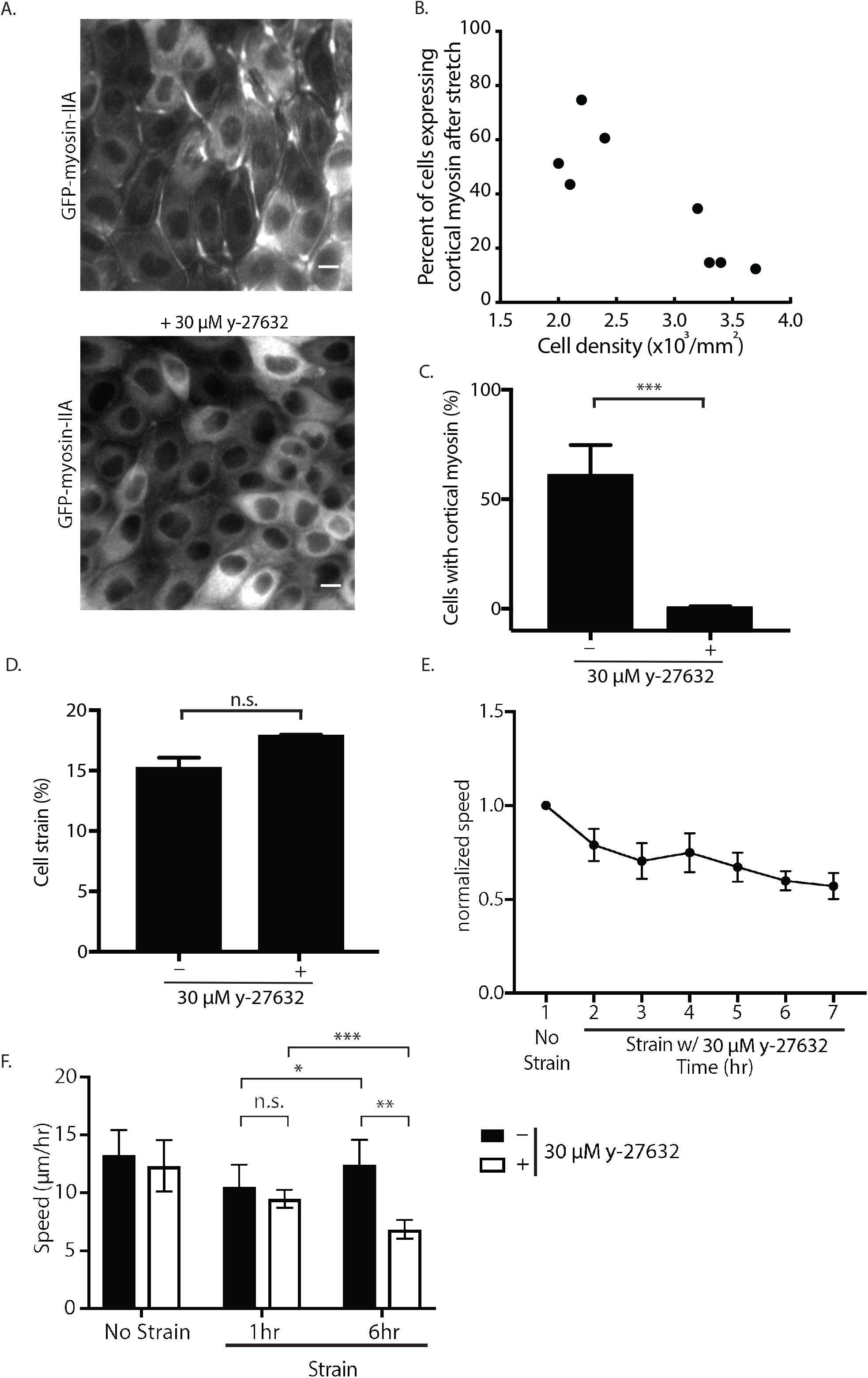
Dynamics of collective cell migration in response to stretch is dependent on cell density and myosin activation downstream of Rho-associated protein kinase (ROCK). A. Localization of GFP-Myosin-IIA in MDCK cells that were plated for 20 hr in the cell stretching device and stretched at 15% for 1 hr with or without the ROCK inhibitor (30 μM y-27632) added immediately before the application of stretch. Scale bars: 10 μm. B. Quantification of the number of cells expressing cortical myosin after the application of 15% stretch for 1 hr with varying cell density from 2 × 10^3^/mm^2^ to 3.7 × 10^3^/mm^2^. Quantifications were from 8 independent experiments with at least 900 cells per experiment. C. Quantification of the number of cells expressing cortical myosin after the application of 15% stretch with or without the ROCK inhibitor (30 μM y-27632). Quantifications were from 3 independent experiments with at least 900 cells analyzed per experiment. D. Quantifications were mean +/− SD; unpaired t-test p values; ***, p<0.001. Quantification of the individual cell strains at 35 kPa of vacuum pressure with or without the ROCK inhibitor (30 μM y-27632). Quantifications were mean +/− SD from 3 independent experiments. E. Normalized mean velocities over the course of 7 hr of imaging every 10 min with strain and the ROCK inhibitor (30 μM y-27632). Hours 2–7 were normalized to the first hour of no strain and no inhibitor for all experiments. Quantifications were mean +/− SD from 4 independent experiments. F. Mean velocities of MDCK cells treated with or without ROCK inhibitor (30 μM y-27632) at 1 hr or 6 hr of strain compared to those under no strain. Quantifications were mean +/− SD from 4 independent experiments; paired t-test p-value for comparison between 1 hour and 6 hours for the same monolayer; unpaired t-test p-value for comparison of monolayers treated with or without ROCK inhibitor (30 μM y-27632); ***, p<0.001; **, p<0.01; *, p<0.05.

### Rho/ROCK activity regulates increases in cell movements

Since applying mechanical stretch to monolayers caused the relocalization of myosin IIA from the cytoplasm to the plasma membrane, we hypothesized that the change in average cell velocity upon stretch (see Figure 4C) was dependent on myosin IIA activity. To test this possibility, we used y-27632 to inhibit the activity of ROCK, which is a positive regulator of myosin IIA tension.^31,32^ Addition of 30 μM y-27632 upon application of stretch resulted in little or no accumulation of GFP-myosin IIA at the cell cortex in cells under stretch [0.73 ± 0.34% (mean ± SD); Figure 5A, C] compared to cells stretched in the absence of y-27632 [62.2 ± 11.8% (mean ± SD); Figure 5A, C]. Note that our CSI code showed that there was no significant difference in cell strain in the presence or absence of the ROCK inhibitor at 35 kPa (Figure 5D). PIV analysis showed that the average cell velocity in y-27632-treated cell monolayers decreased with (Figure 5E) or without mechanical stretch (Figure S5B). Incubation of cells with y-27632 resulted in a decrease in cell velocity over 6 hr compared to cells under stretch in the absence of the inhibitor (Figure 5F), indicating that the recovery of cell velocity measured in stretched monolayers (Figure 4C) was dependent on ROCK activity.

## Discussion

This study describes the design and fabrication of a device that applies a constant, uniaxial stretch using materials, reagents and equipment readily available to most laboratories. This device has advantages over others because it is relatively inexpensive to fabricate from a 3D-printed mold and PDMS, and has a simple, vacuum-based design (Figures 1, 2).

We developed MATLAB code (CSI) that accurately measures individual cell strains, which match the predicted strain applied to the device (Figures 1 and 3). These measured cell strains are consistent with a previous study of the strain profile for this device design made by tracking the displacement of fluorescent beads added to the surface of the substrate.^13^ Measuring cell strains compared to bead displacement gave insight into the heterogeneity of cell strains across a monolayer. The heterogeneity measured by our code could be explained by differences in local cell density within the monolayer or cell shape differences. Our CSI code also showed that cell areas changed at a rate similar to that of cell strains, but the cell perimeter had smaller relative changes with increasing vacuum pressure. This could be explained by changes in the perimeter occurring at a different z-plane from the imaging z-plane, possibly at the basal membrane of the cell (Figure 3).

As an application, we examined: 1) the collective cell behavior in an epithelial monolayer under stretch; and 2) the role of actomyosin tension by imaging the distribution of GFP-myosin IIA and the effects of addition of the ROCK inhibitor y-27632. We found that the overall speed of individual cell movements within the cell monolayer decreased after stretch and that this response was dependent on cell density of the monolayer. High-density cell monolayers did not accumulate cortical myosin IIA and did not respond by slowing down in response to stretch (Figure 4). These results demonstrate that cell density is a strong regulator of cell movements in response to mechanical stretch.

We detected a significant increase in the level of cortical myosin IIA during application of strain that was dependent on ROCK activity (Figure 5). Significantly, after 2 hr of strain the velocity of cells under stretch began to increase to a level similar to that before the application of stretch (while stretch was maintained for 7 hr). This increase in cell movements required intact ROCK activity (Figure 5). In contrast, the velocity of cells not under strain gradually decreased over 7 hr. A parsimonious explanation of these data is that mechanical stretch activates ROCK,^19^ which induces the cortical accumulation of myosin IIA,^1,2^ possibly through changes in tension at cell-cell contacts.^4^ An increase in cortical tension leads to decreases in cell movement and rearrangements.^5^ However, we observed an initial decrease in cell movements that is independent of the accumulation myosin IIA at cell-cell contacts suggesting that effects of strain on ROCK activity at cell adhesions to the extracellular matrix (ECM) may be important (Figure 5).^24^ This is supported by recent work showing that traction forces at cell-ECM junctions, and not cortical tension at cell-cell junctions, regulate cell movements and rearrangements.^33^ These data indicate that cortical myosin IIA localization and ROCK activity do not regulate the initial slowing of cell movements after stretch, but regulate the increase in cell speeds over longer times in response to mechanical stretch.

Increased cell density has been shown to lead to a transition from a fluid state to a more solid, glassy state.^3^ Our results indicate that mechanical stretch can cause a low-density monolayer, which is in a more fluid state, to transition to a more solid state and behave like a high-density monolayer as evidenced by the decrease in cell movements. Since high-density monolayers are in a more solid state, they do not undergo this transition upon mechanical stretch. Furthermore, over the course of 7 hr under stretch we found that cell movement returns to levels observed before stretch. Thus, a low-density cell monolayer under stretch transitions to a more solid state, but then returns to a more fluid state in a process that is dependent on Rho/ROCK activity. This transition from fluid to solid states could be a cellular response to withstand the increase in mechanical force on the monolayer under stretch. This process may contribute to the overall maintenance of epithelial homeostasis,^27^ and indicates that epithelial tissue can respond to a mechanical force with material-like transitions between fluid and solid states.

In summary, we have validated the usefulness of the device, system, and methods described here by examining the effects of stretch on collective cell movement, and the role of cell density and actomyosin tension in mediating effects of strain. A prototype of the device was also instrumental in studying how spindle orientation is affected by the direction of uniaxial mechanical stretch.^13^ The detailed methods provided here for the fabrication, use of the device, and the software to analyze changes in cell properties at the single-cell level will make mechanobiology studies more accessible to a broader range of bioengineers and biologists.

## Acknowledgments

We thank So Yamada (University of California, Davis) for the MDCK GFP-myosin IIA cells. This work was supported by a NIH Training Grant T32GM007276 (to K.C.H.), a National Science Foundation (NSF) Graduate Research Fellowships Program (to J.T.), a Life Science Research Foundation Fellowship (Howard Hughes Medical Institute sponsorship) (to D.J.C.), NSF grants (CMMI 1662431 and EFRI MIKS 1136790 to W.J.N. and B.L.P.), a Stanford Bio-X research fellowship (to K.C.H., J.T., and J.Y.S.), a Stanford Bio-X grant (to W.J.N. and B.L.P.), the Ilju Foundation (J.Y.S.), and NIH Grant R35GM118064 (to W.J.N.).

## Supplemental Figures

**Supplemental Figure 1:**
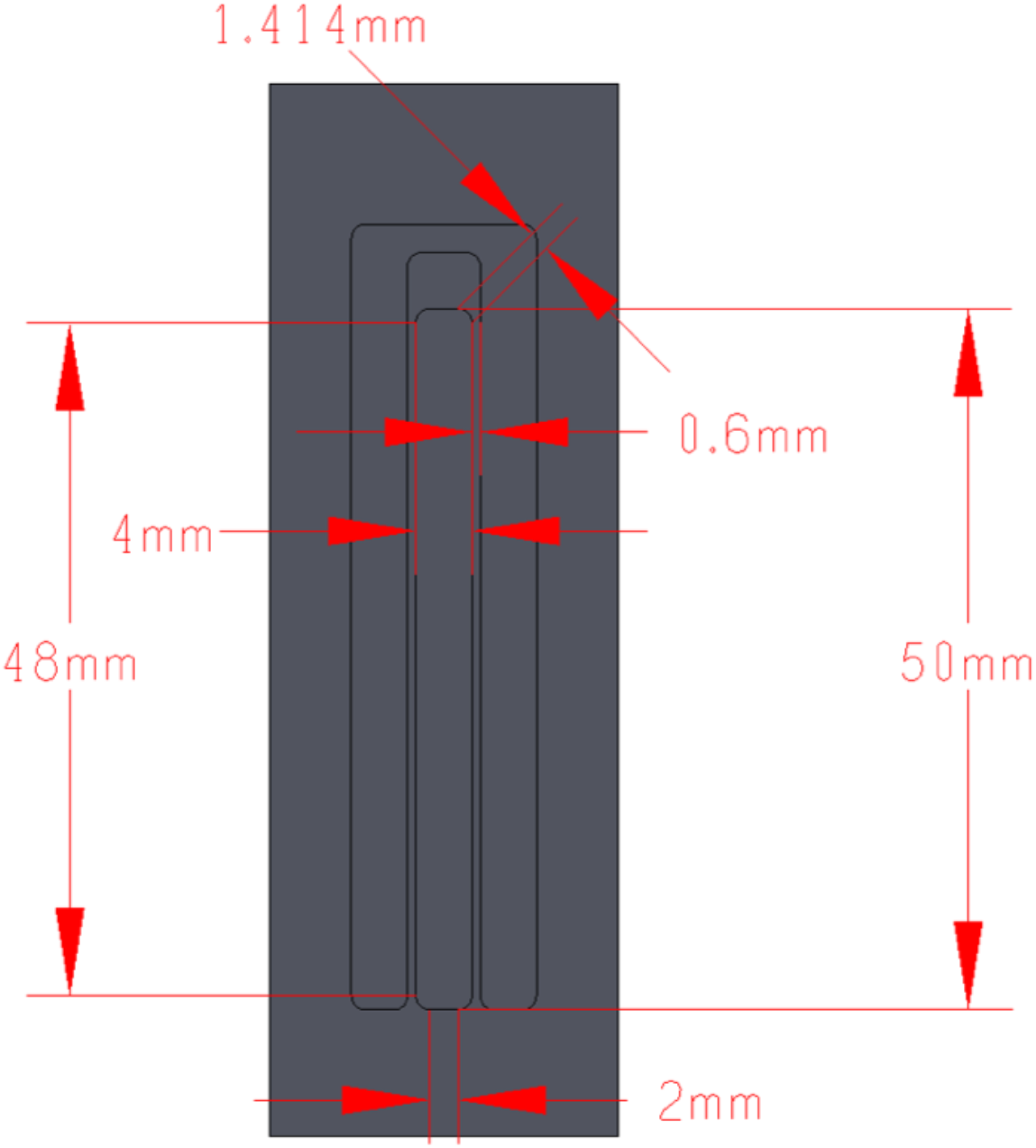
Dimensions of the 3-D printed mold of the cell stretching device.

**Supplemental Figure 2:**
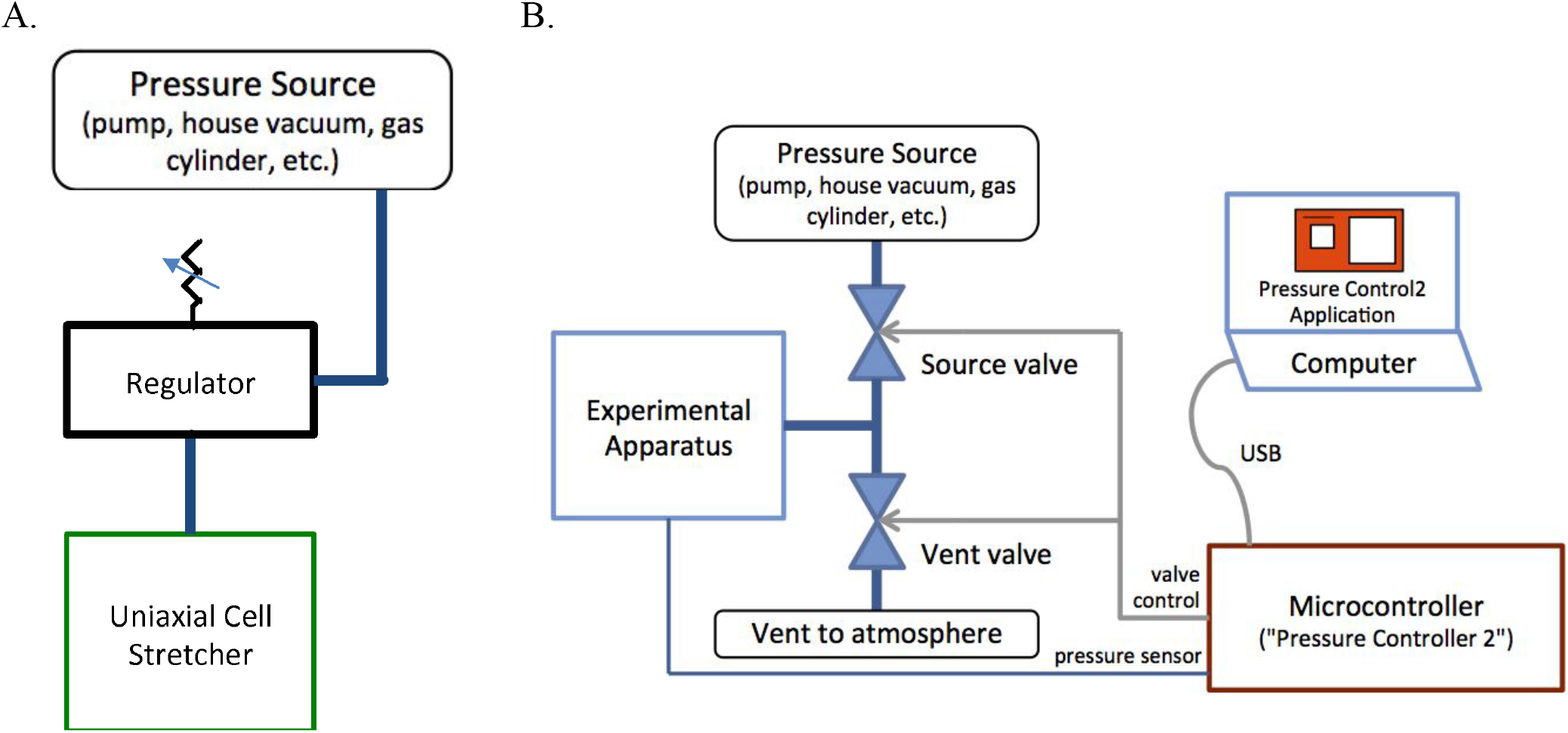
Schematic diagram of the two types of pressure regulators used in the experiments. A. Schematic diagram of the mechanical regulator, High Precision Vacuum Regulator-P3RA171, from Parker. The vacuum regulator connects the in-house vacuum line, with a vacuum pressure of 90 kPa, to the device. B. Schematic diagram of the electronic pressure controller from Red Dog Research. This electronic controller connects the in-house vacuum line to a valve controller that is connected to a laptop. The application on the computer can control the valves to within 0.25 kPa of the setpoint and it has the capacity to execute pre-programmed pressure waveforms, and hence modulate strain in a variety of ways, including sinusoid waveforms.

**Supplemental Figure 3:**
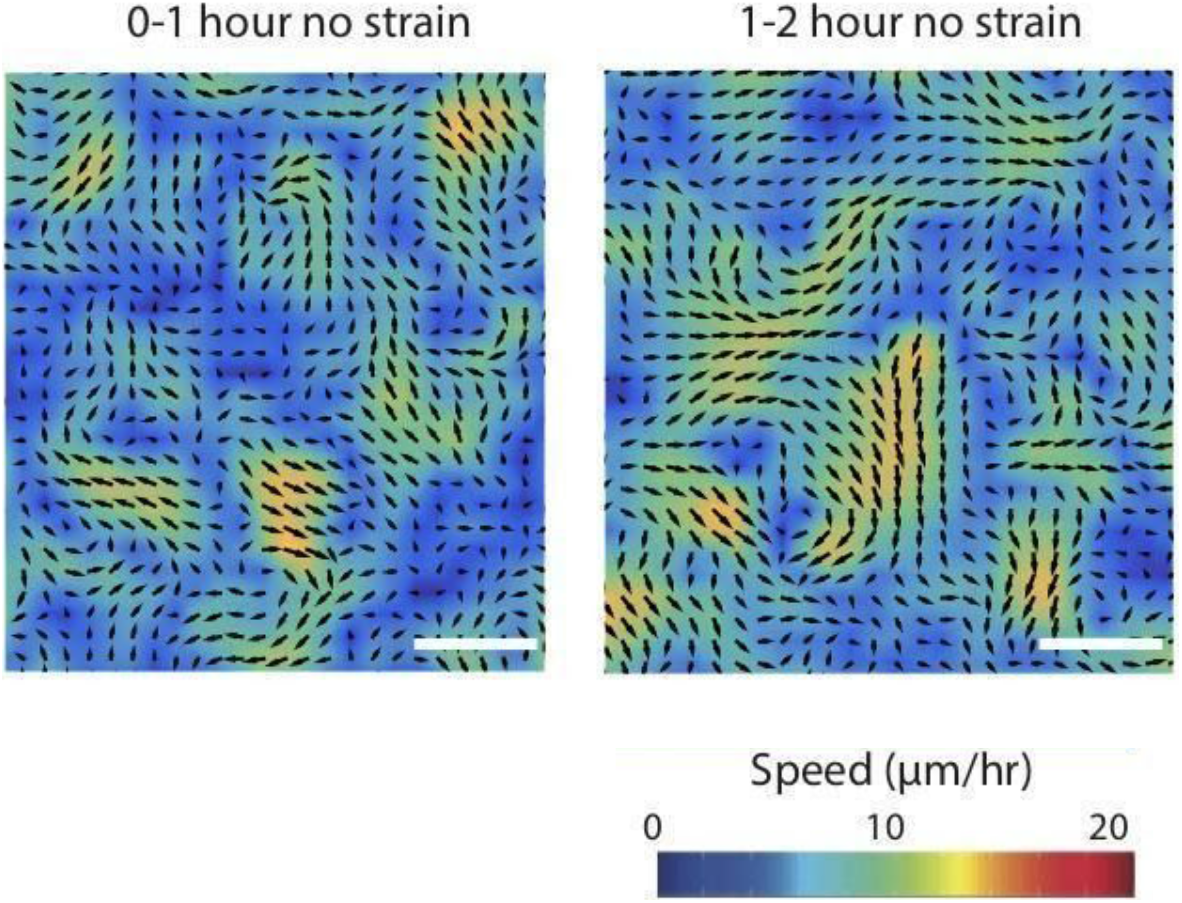
Mean velocities of MDCK cell monolayers under no strain conditions over the course of 2 hr. Cell monolayers were imaged with phase contrast every 10 min at 5x magnification. The cell monolayer was imaged for 1 hr followed by another hour with no strain. The collective migration velocities were measured with PIV (particle image velocimetry). Scale bars: 50 μm.

**Supplemental Figure 4:**
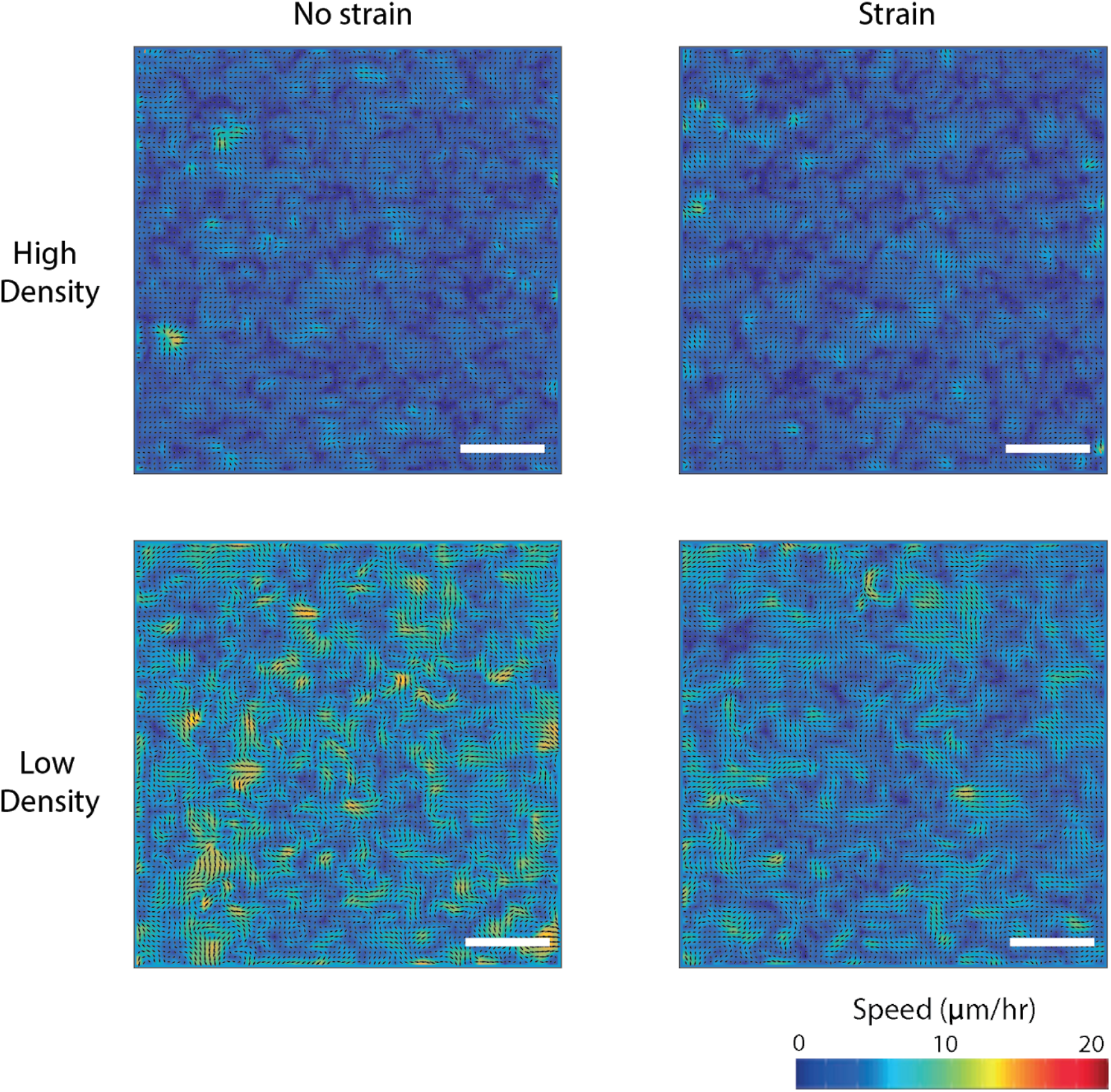
Representative mean velocity maps from 1 hr of imaging, calculated from the phase contrast images taken every 10 min at 5× magnification of both low-density (2.0 ×10^3^/mm^2^ –2.5 ×10^3^/mm^2^) and high-density (3.0 ×10^3^/mm^2^ –4.0 ×10^3^/mm^2^) monolayers. The cell monolayers were imaged for 1 hr followed by another hour with or without strain. The collective migration velocities were measured with PIV (particle image velocimetry). Scale bars: 150 μm.

**Supplemental Figure 5:**
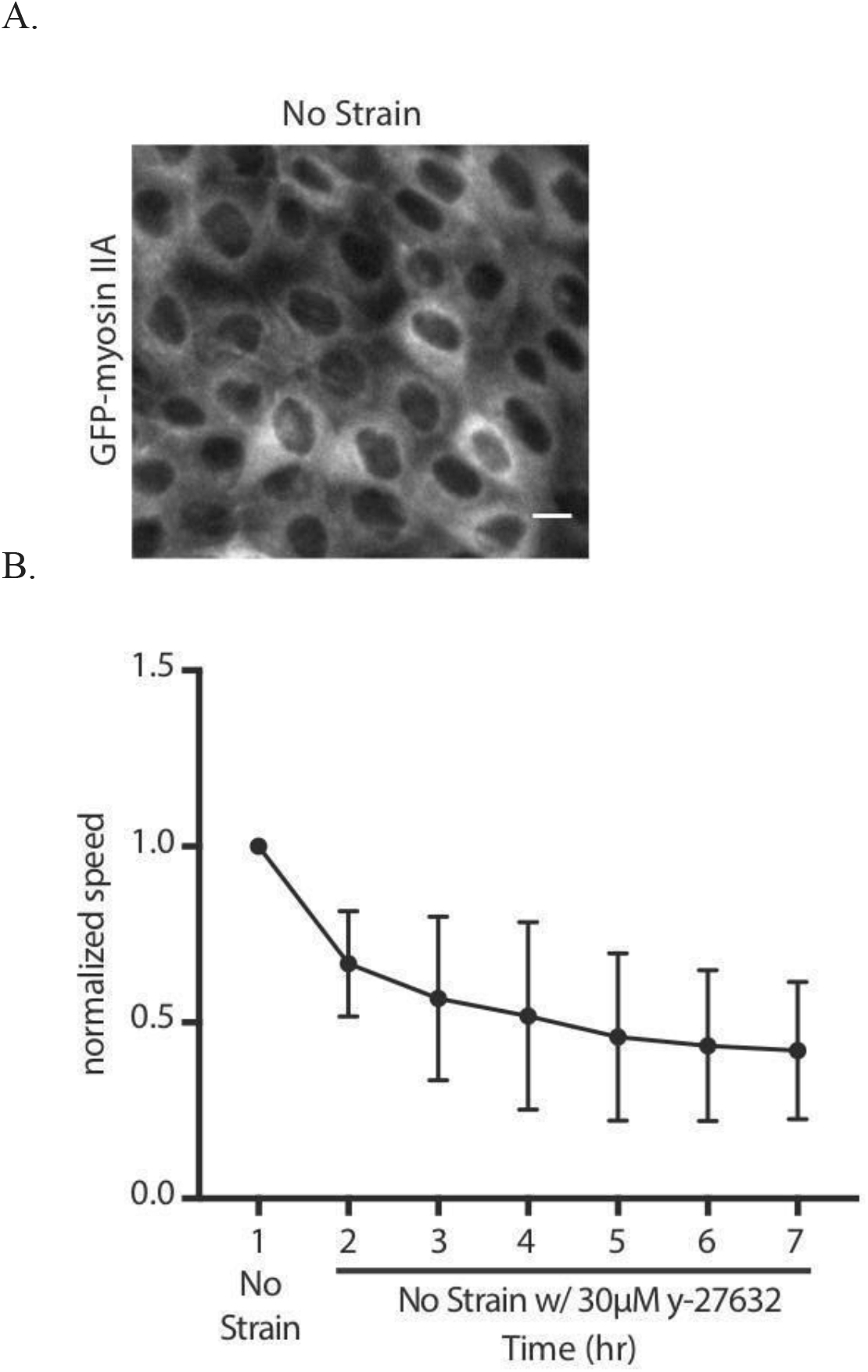
A. Representative image of MDCK cells expressing GFP-Myosin IIA under no strain where we do not see the accumulation of GFP myosin at the cortex of cells. Scale bars: 10μm B. Normalized mean velocities over the course of 7 hr of imaging every 10 minutes with no strain and the ROCK inhibitor (30 μM y-27632). Hours 2–7 were normalized to the first hour of no strain and no inhibitor for all experiments. Quantifications were mean +/− SD from 3 independent experiments.

**Supplemental Figure 6:**
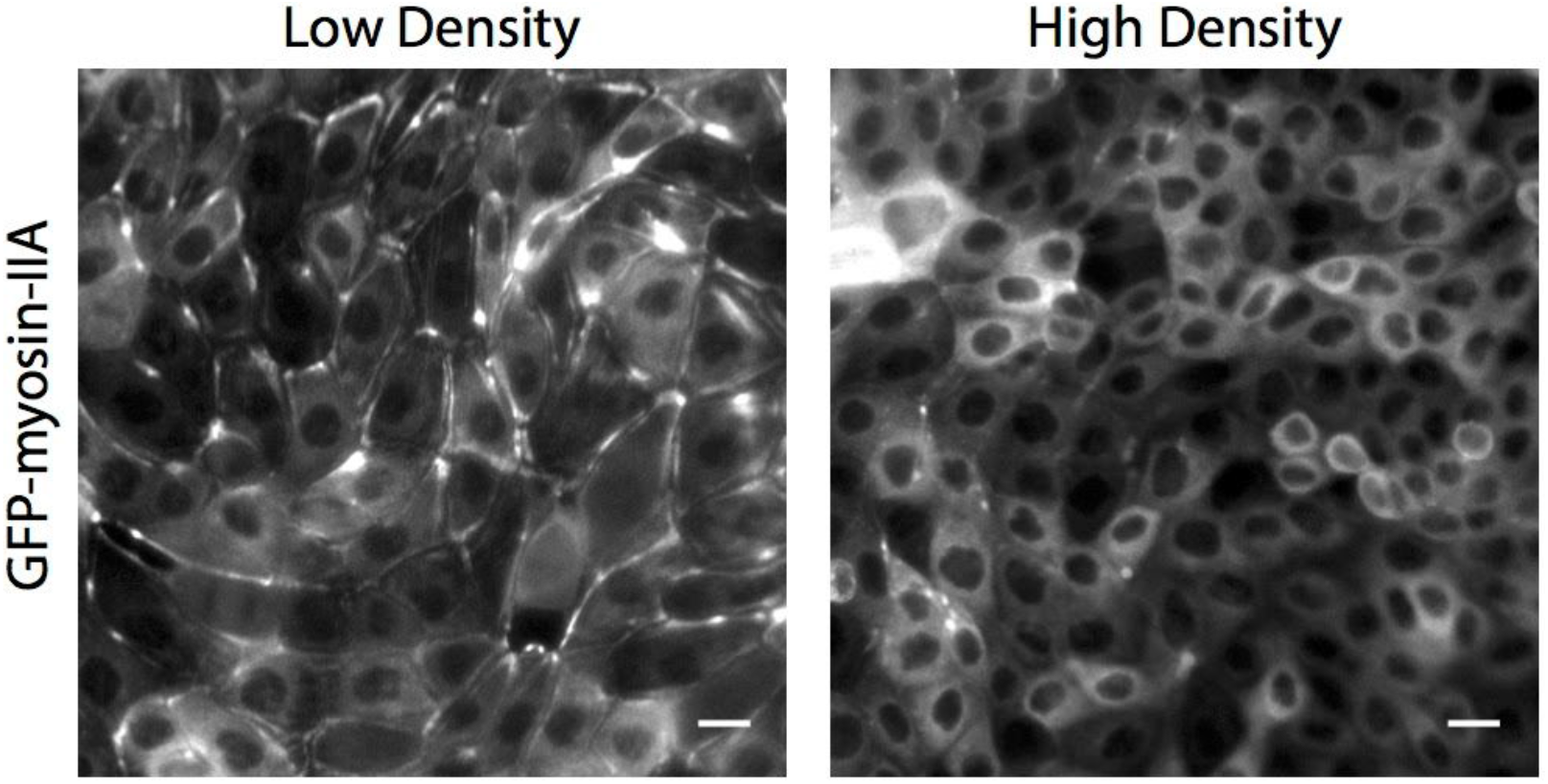
Representative image of MDCK cells expressing GFP-Myosin IIA grown at low and high density under 15% strain for 1 hr. Scale bars: 10μm

**Supplemental Figure 7:**
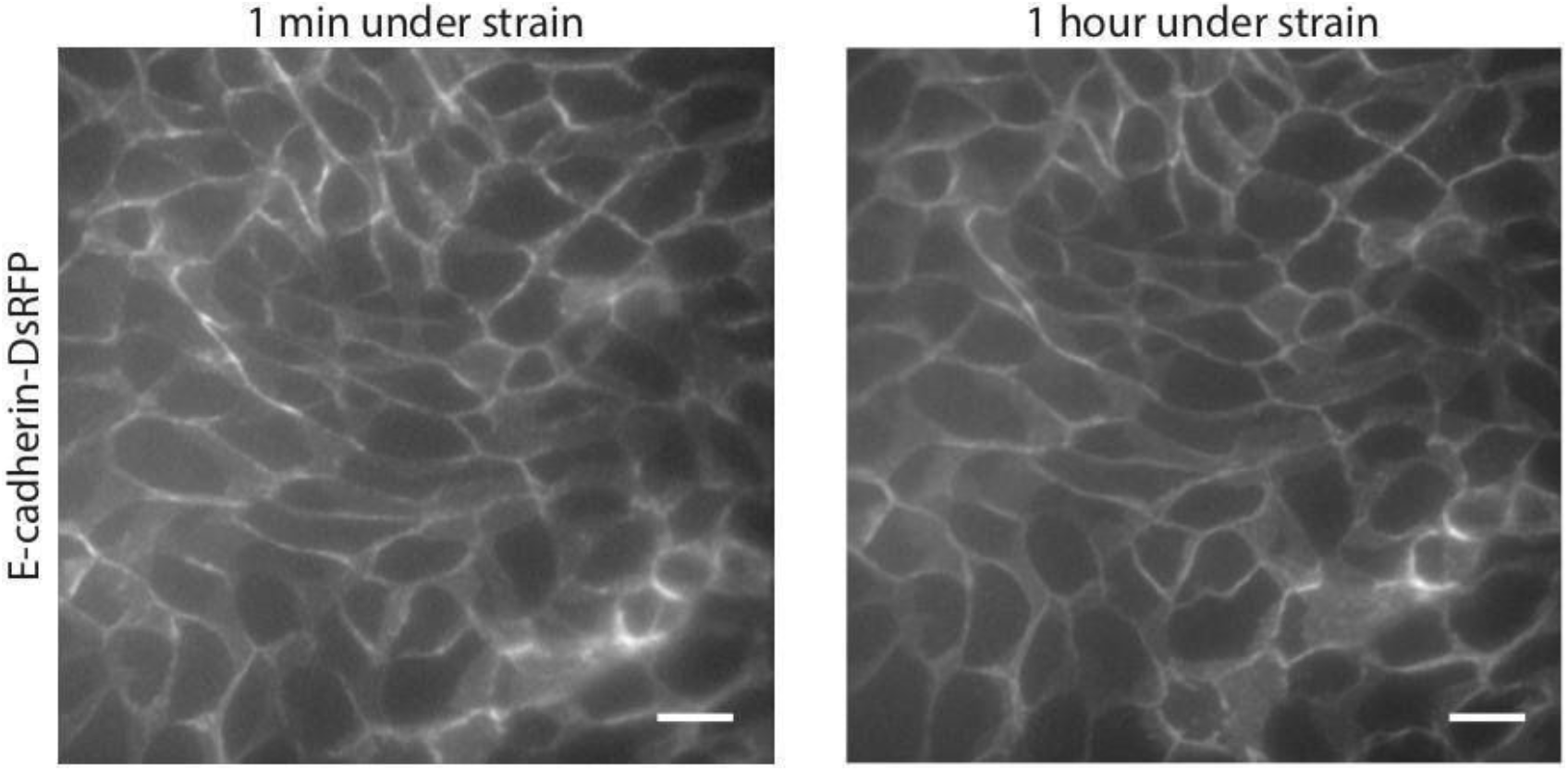
Representative images of MDCK cells expressing E-cadherin-DsRFP under 1 min and 1 hr of strain where we see no changes in E-cadherin. Scale bars: 10μm.

**Supplemental Movie 1:**
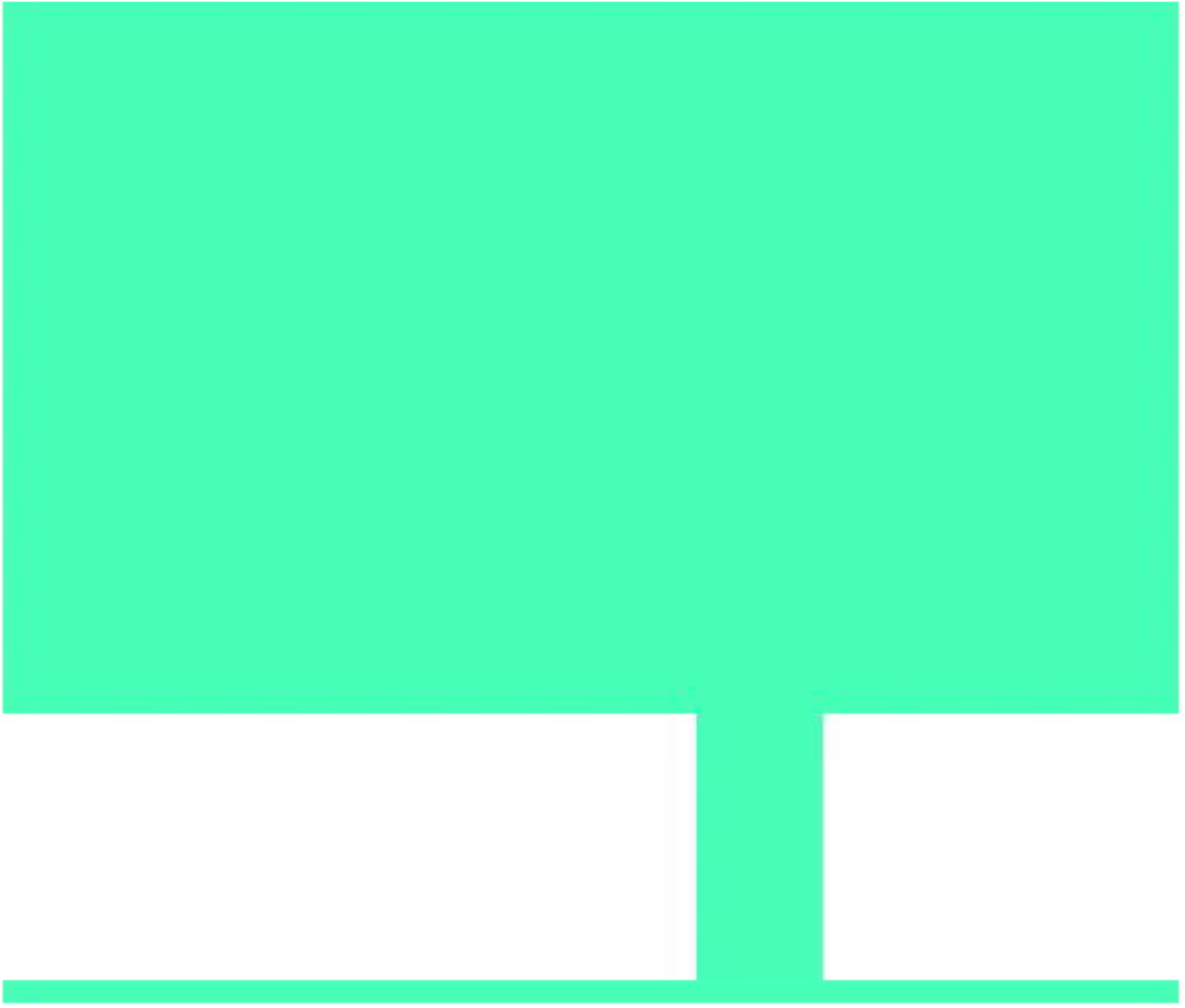
2D FEA simulation of the strain applied to the cell culture membrane from 0 kPa to 70 kPa of vacuum pressure to the side chambers using COMSOL Multiphysics (version 4.4, COMSOL Inc.).

**Supplemental Movie 2:**
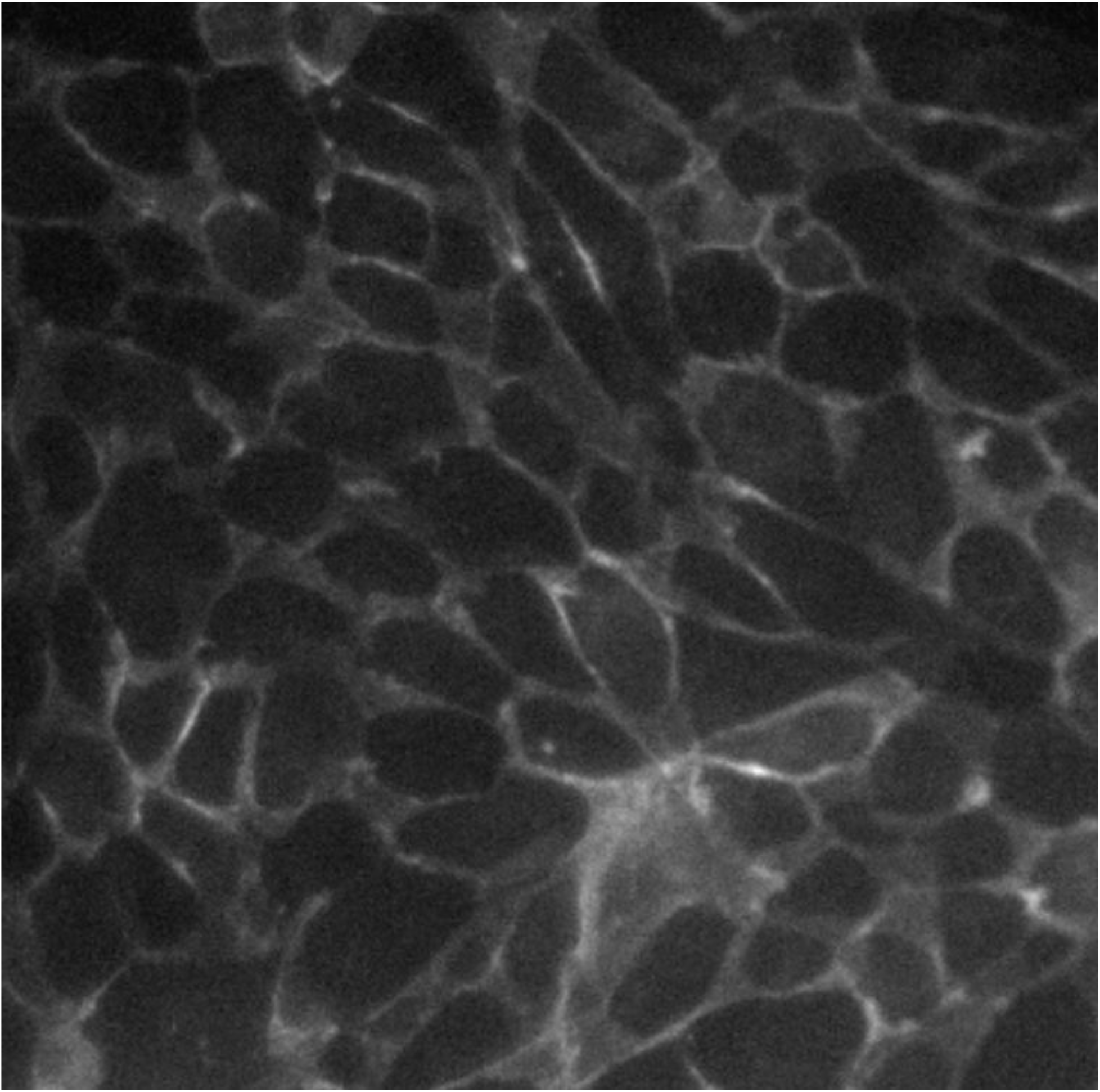
E-cadherin-RFP MDCK cells plated 20 hr before imaging on the cell stretching device. Image acquired every 3.5 kPa from 0 kPa to 70 kPa at 40x magnification at 5-sec interval between images.

